# Beyond massive univariate tests: Covariance regression reveals complex patterns of functional connectivity related to attention-deficit/hyperactivity disorder, age, sex, and response control

**DOI:** 10.1101/2021.02.09.430522

**Authors:** Yi Zhao, Mary Beth Nebel, Brian S. Caffo, Stewart H. Mostofsky, Keri S. Rosch

**Affiliations:** Department of Biostatistics and Health Data Science, Indiana University School of Medicine, Indianapolis, IN, USA; Center for Neurodevelopmental and Imaging Research, Kennedy Krieger Institute, Baltimore, MD, USA; Department of Neurology, Johns Hopkins University School of Medicine, Baltimore, MD, USA; Department of Biostatistics, Johns Hopkins University Bloomberg School of Public Health, Baltimore, MD USA; Department of Psychiatry and Behavioral Sciences, Johns Hopkins University School of Medicine, Baltimore, MD USA; Department of Neuropsychology, Kennedy Krieger Institute, Baltimore, MD, USA

**Keywords:** functional connectivity, ADHD, response control, covariance regression, sex differences, childre

## Abstract

We applied a novel Covariate Assisted Principal (CAP) whole-matrix regression approach to identify resting-state functional connectivity (FC) brain networks associated with attention-deficit/hyperactivity disorder (ADHD) and response control. Participants included 8-12 year-old children with ADHD (n=115, 29 girls) and typically developing controls (n=102, 35 girls) with a resting-state fMRI scan and go/no-go task behavioral data. We modeled three sets of covariates to identify resting-state networks associated with ADHD, age, sex, and response control. Four networks were identified across models revealing complex interactions between subregions of cognitive control, default mode, subcortical, visual, and somatomotor networks that relate to age, response control, and a diagnosis of ADHD among girls and boys. Unique networks were also identified in each of the three models suggesting some specificity to the covariates of interest. These findings demonstrate the utility of our novel covariance regression approach to studying functional brain networks relevant for development, behavior, and psychopathology.

## Introduction

Attention-deficit/hyperactivity disorder (ADHD) is a common neurodevelopmental disorder that affects approximately 5-10% of children and adolescents.^1^ ADHD is characterized by developmentally inappropriate symptoms of inattention and/or hyperactivity/impulsivity that impact academic, family, and social functioning.^2^ The diagnosis of ADHD is based on parent and teacher report of the behavioral symptoms of ADHD. These behavioral symptoms are thought to arise from cognitive, motor, and motivational deficits and associated atypical brain structure and function. In particular, impaired response control, including poor inhibition and increased trial-to-trial variability, are among the most consistent findings in the ADHD neurocognitive literature and central to theoretical models of ADHD.^3,4^ Within the ADHD literature, there is an increased focus on elucidating the neurobiological basis for ADHD to impaired response control to ultimately inform prevention and intervention for this highly prevalent and impairing disorder.

Neuroimaging methods have increasingly been applied to characterize atypical brain structure and function in individuals with ADHD. In particular, functional MRI during a “resting state” (rs-fMRI) is widely being used to examine functional networks that operate differently in children with ADHD and relate to neurocognitive deficits associated with the disorder. The extant literature has shown that the neurobiological basis for ADHD likely involves dysfunctional interactions of, or functional connectivity (FC) between, brain networks rather than atypical structure or function of isolated brain regions. Default mode network (DMN) hyperconnectivity with other networks among individuals with ADHD, which reflects increased integration between said networks (and reduced modularity), is one of the most consistent findings in the literature.^5–9^ Some studies have also reported DMN hyperconnectivity with other networks to relate to cognitive deficits in individuals with ADHD.^6,10,11^ However, the ADHD rs-FC literature extends beyond DMN connectivity to include findings of increased within-network connectivity in motor, ^12–14^ and visual regions,^9,13,15,16^ and atypical FC between frontal-subcortical regions.^17–20^ Thus, a whole-brain approach is warranted to better characterize differences in functional network organization in ADHD. Moving beyond diagnostic group comparisons, characterization of how individual differences in functional networks are associated with behavior has the potential to inform the heterogeneity of neurocognitive deficits and symptom presentation in children diagnosed with ADHD.

A typical strategy of studying brain functional connectivity is to perform statistical analysis on each individual connection. One drawback of this element-wise approach is multiplicity. Given a set of *p* brain voxels/regions, statistical inference needs to account for at least *p*(*p* − 1)/2 hypothesis tests, one for each element of the connectivity matrix. To circumvent this multiplicity issue, Zhao et al.^21^ proposed a whole-matrix regression approach referred to as Covariate Assisted Principal (CAP) regression. It aims to identify a common linear projection of *p* time courses across subjects such that variations in functional connectivity defined by the projection can be explained by the covariates of interest. It is a meso-scale approach in the sense that, with an appropriate threshold, the projection defines a brain subnetwork. However, this approach suffers from the “curse of dimensionality”, in that the dimension of the data, *p*, cannot be greater than the number of fMRI volumes. Therefore, it cannot be applied to voxel-level fMRI data. Building off the CAP regression approach, an integrative approach was proposed to analyze whole brain voxel-level data, revealing individual and group variations in functional connectivity.^22^ This integrative approach starts with a dimension reduction step, such as group independent component analysis (ICA),^23^ which is followed by CAP regression on the ICs. Projecting back to the voxel space, it yields a reconstructed brain map that is associated with the covariates.

The current study applies this novel integrated CAP analysis to examine how a diagnosis of ADHD, relevant demographic variables, and behavioral measures of response control predict functional interactions between brain networks. Specifically, we applied this novel, high-dimensional statistical method to study interactions between sex and an ADHD diagnosis on FC in children with and without ADHD and in relation to cognitive deficits associated with ADHD.

## Results

The CAP method identified five brain subnetworks (groups of RSN ICs, or mega-component) in the diagnosis-sex model, six in the GNG tau model, and seven in the GNG CR model. We summarize the significance of the coefficient effect in each mega-component in **Table 1**. **Supplementary Figure S1** summarizes the RSN ICs clustered within the functional groups described above that contribute to each component as river plots. Among all of these mega-components, four subnetworks were identified in all three models. **Figure 1** compares the similarity between the mega-components identified by the three models; a connection between two mega-components suggests a high similarity. **Figure 2** illustrates the loadings of RSNs contributing to each mega-component as reconstructed brain maps (**Supplemental File S1** is an interactive tool to view 3D images of the ten mega-components to accompany Figure 2a-j).

**Table 1.**
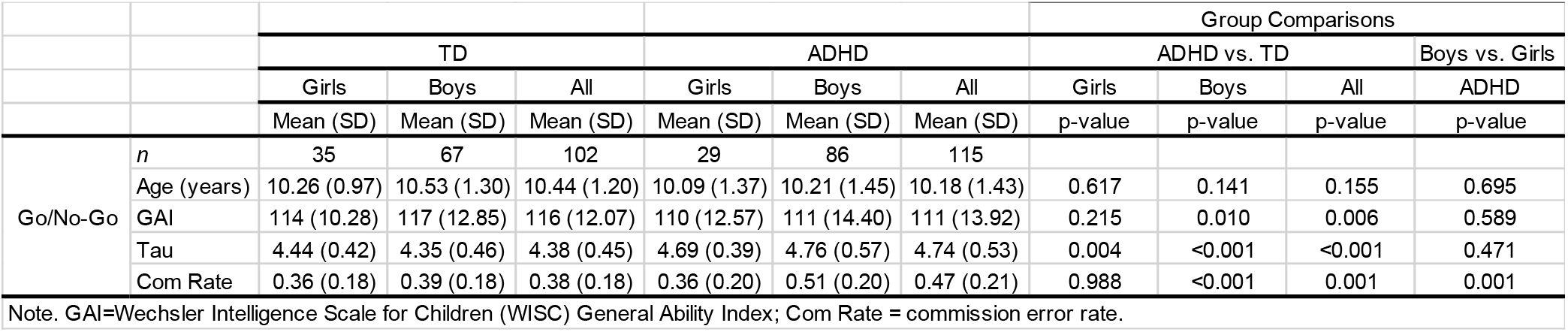
Behavioral task performance and relevant demographics for the ADHD and TD groups overall and within sex.

|  |  | TD |  |  | ADHD |  |  | Group Comparisons |  |  |  |
| --- | --- | --- | --- | --- | --- | --- | --- | --- | --- | --- | --- |
|  |  |  |  |  |  |  |  | ADHD vs. TD |  |  | Boys vs. Girls |
|  |  | Girls | Boys | All | Girls | Boys | All | Girls | Boys | All | ADHD |
|  |  | Mean (SD) | Mean (SD) | Mean (SD) | Mean (SD) | Mean (SD) | Mean (SD) | p-value | p-value | p-value | p-value |
| Go/No-Go | *n* | 35 | 67 | 102 | 29 | 86 | 115 |  |  |  |  |
|  | Age (years) | 10.26 (0.97) | 10.53 (1.30) | 10.44 (1.20) | 10.09 (1.37) | 10.21 (1.45) | 10.18 (1.43) | 0.617 | 0.141 | 0.155 | 0.695 |
|  | GAI | 114 (10.28) | 117 (12.85) | 116 (12.07) | 110 (12.57) | 111 (14.40) | 111 (13.92) | 0.215 | 0.010 | 0.006 | 0.589 |
|  | Tau | 4.44 (0.42) | 4.35 (0.46) | 4.38 (0.45) | 4.69 (0.39) | 4.76 (0.57) | 4.74 (0.53) | 0.004 | <0.001 | <0.001 | 0.471 |
|  | Com Rate | 0.36 (0.18) | 0.39 (0.18) | 0.38 (0.18) | 0.36 (0.20) | 0.51 (0.20) | 0.47 (0.21) | 0.988 | <0.001 | 0.001 | 0.001 |

Note. GAI=Wechsler Intelligence Scale for Children (WISC) General Ability Index; Com Rate = commission error rate.

**Figure 1.**
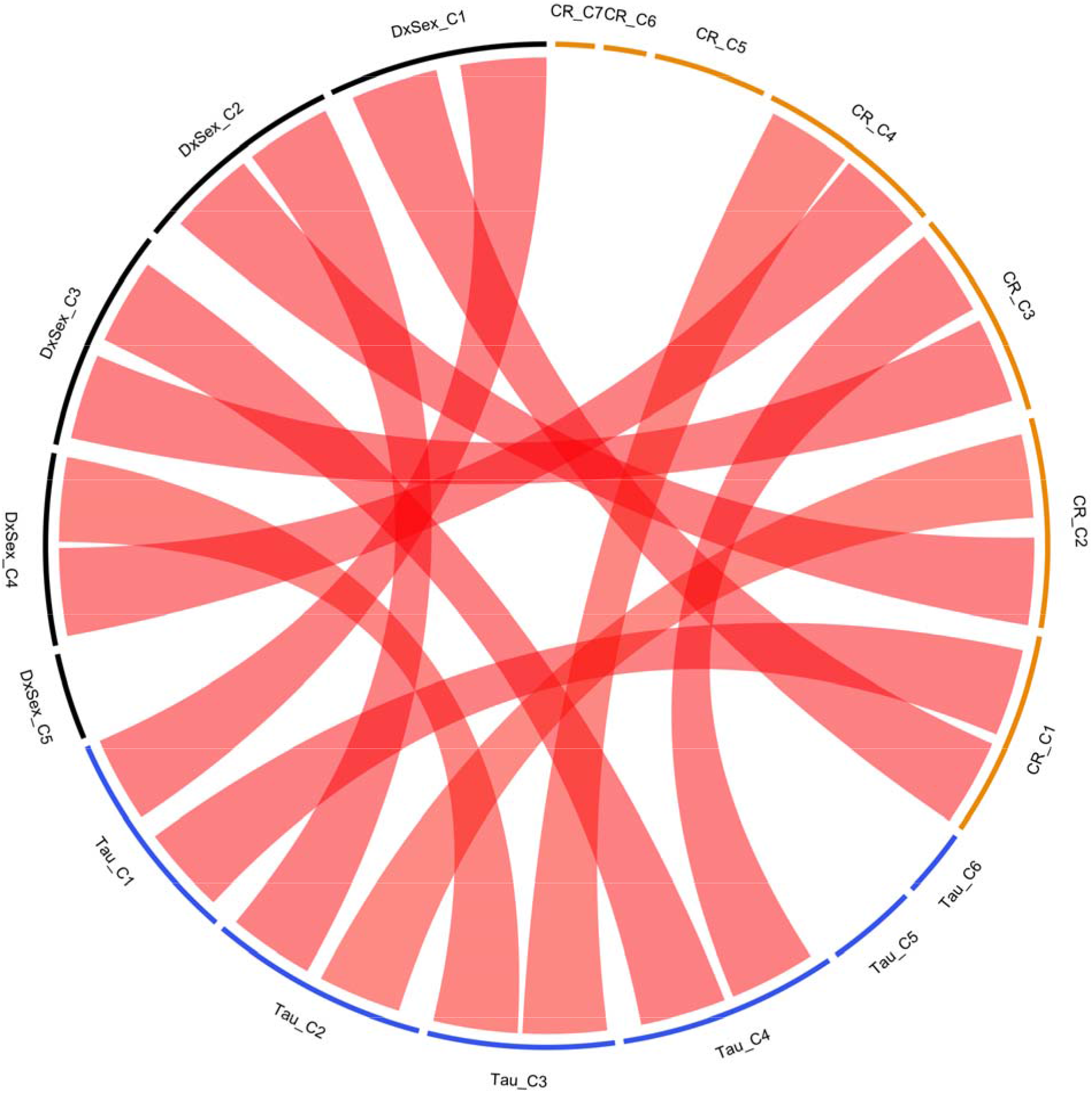
Chord diagram compares the similarity between the components identified from the three models. DxSex: the behavior free model; Tau: model including Go/No-Go Tau as one of the predictors; CR: model including Go/No-Go commission error rate as one of the predictors. A red connection indicates that the two components are highly similar.

**Figure 2.**
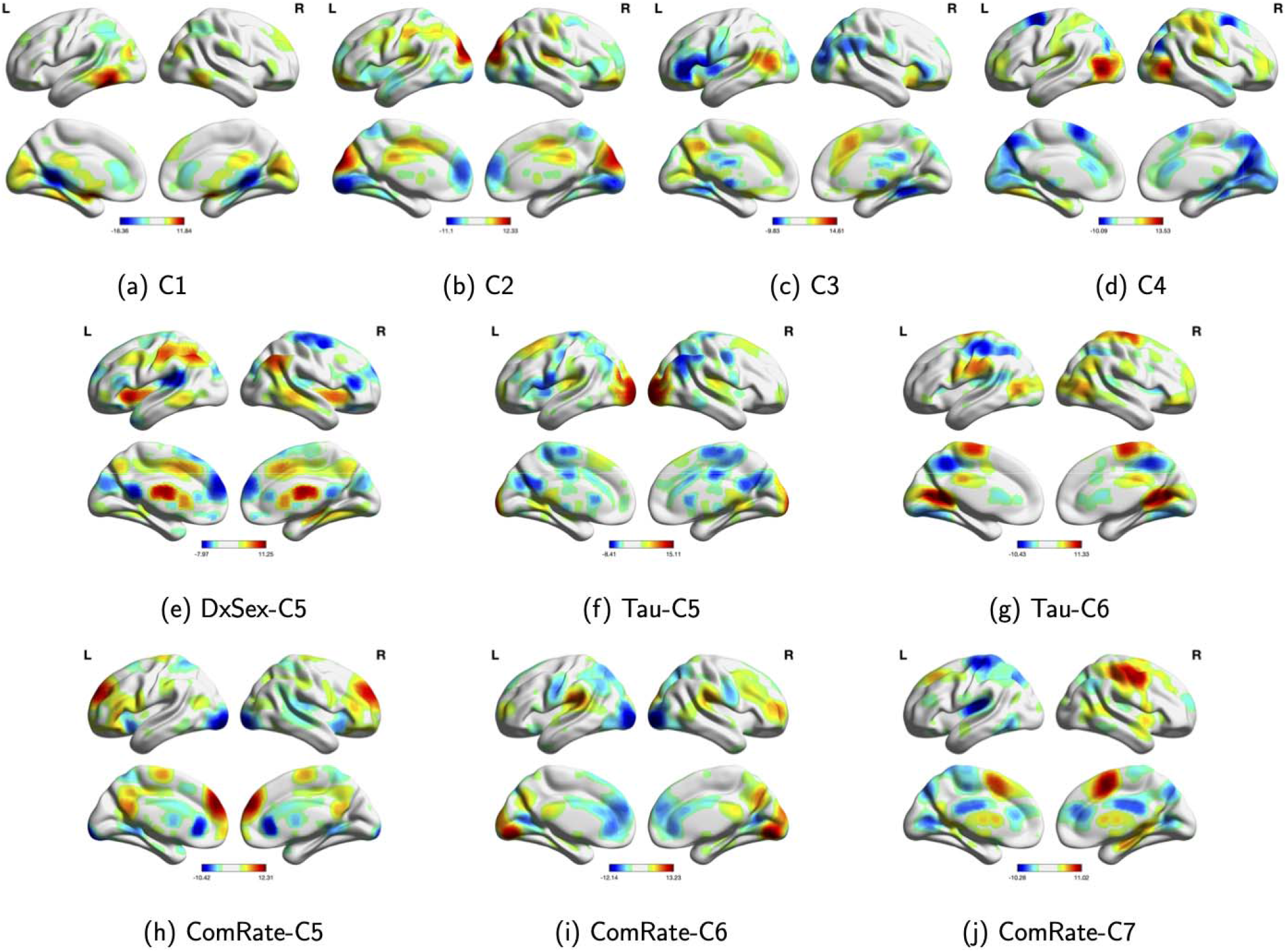
Reconstructed brain maps of the components identified by the CAP method. DxSex: the behavior free model; Tau: model including Go/No-Go Tau as one of the predictors; ComRate: model including Go/No-Go commission error rate as one of the predictors. (a)-(d) Common components identified by all three models. (e) Unique component identified by the behavior-free model. (f)-(g) Unique components identified by the model with Go/No-Go Tau as one of the predictors. (h)-(j) Unique components identified by the model with Go/No-Go commission error rate as one of the predictors.

### First Common Mega-Component (DxSex-C1/Tau-C1/CR-C1; *Figure 2a*)

Across the three models, the identified FC mega-component consisted of subregions of the cognitive control (inferior temporal gyrus; ITG) and subcortical (striatal) networks with positive loadings, as well as subregions of the DMN (posterior cingulate cortex; PCC) and subcortical (thalamus) networks with negative loadings. This mega-component was positively related to age in each of the models (all *p*s<.001), such that as age increases, FC between regions with the same loading signs increases (ITG-striatal FC and PCC-thalamus FC), while FC between regions with opposite loading signs decreases (e.g., PCC-striatum).

### Second Common Mega-Component (DxSex-C2/Tau-C2/CR-C2; *Figure 2b*)

The second component identified across all three models consisted of subregions of the cognitive control (left central executive), visual (lateral occipital cortex), and sensorimotor (ACC/BA6) networks contributing positively, as well as other subregions of the cognitive control (insula, ITG) and DMN (PCC) networks contributing negatively. This component was negatively associated with GAI (DxSex C2 *p*=.014; Tau C2 *p*=.039; CR C2 *p*=.042) and positively associated with Tau in boys with ADHD (*p*=.001), such that lower GAI and greater Tau is associated with greater FC between regions with the same loading sign and lower FC between regions with opposite loading signs. Girls with ADHD showed greater FC within this mega-component compared to TD girls (DxSex C2, *p*=.022) and lower FC to boys with ADHD (DxSex C2, *p*<.001). However, this diagnostic difference was not observed when controlling for Tau and CR (see **Table 1**).

### Third Common Mega-Component (DxSex-C3/Tau-C4/CR-C3; *Figure 2c*)

The third component identified across all three models consisted of a DMN subregion (precuneus) with positive loadings, as well as subregions of the cognitive control (inferior frontal gyrus; IFG, insula), DMN (PCC) and visual (lateral occipital cortex) networks with negative loadings. Again, this component was positively associated with age across all three models (all *p*s<.001). In addition, boys with ADHD showed higher FC within this mega-component compared to TD boys (*p*=.009), controlling for age and GAI. However, the diagnostic effect on FC in boys is not significant when controlling for GNG task performance (Tau and CR).

### Fourth Common Mega-Component (DxSex-C4/Tau-C3/CR-C4; *Figure 2d*)

The fourth component identified across all three models consisted of subregions of sensorimotor (right lateralized somatomotor) and visual (lateral occipital cortex) networks with positive loadings, as well as subregions of the DMN (ACC, superior parietal lobe, SMA) and visual network (anterior primary visual; BA17) with negative loadings. This component differed among girls and boys with ADHD (DxSex C4 *p*<.001; Tau C3 *p=*.046; CR C4 *p=*.045), even when controlling for GNG performance, such that boys with ADHD showed stronger FC within this network with and without CR as a covariate. In contrast, girls with ADHD show stronger FC within this network when Tau is included in the model, suggesting network topology differences among girls and boys with ADHD with equivalent response variability. Furthermore, this component was negatively related to Tau (*p*<.001) and CR (*p*=.015) in boys with ADHD, but not among girls with ADHD.

### Unique component in the diagnosis-sex model (DxSex-C5; *Figure 2e*)

This component was only present in the DxSex model, consisting of subregions of the cognitive control (left central executive network, insula), DMN (dlPFC), sensorimotor (caudate, ACC/BA6), and visual (temporal occipital fusiform cortex) and subcortical (caudate) networks with positive loadings, and subregions of the sensorimotor (SMA), visual (visual cortex), DMN (PCC), and auditory (opercular cortex) networks with negative loadings. Results suggest this component is negatively related to age (*p*<.001) and positively related to sex in both ADHD and TD groups, with higher FC in boys (*p*s<.001).

### Unique components in the GNG Tau model (Tau-C5-C6; *Figures 2f-g*)

Two unique components were identified in the GNG Tau model. The first component (Tau C5) consists of subregions of the DMN (PC, SMA), visual (secondary visual area; BA18), and auditory (opercular cortex) networks with positive loadings and subregions of the cognitive control (medial superior frontal cortex), DMN (dlPFC, superior parietal lobe), sensorimotor (ACC/BA6, primary somatosensory, dorsal somatomotor), and subcortical (caudate) networks with negative loadings. Within this component, FC is reduced among girls and boys with ADHD (controlling for age, GAI, and Tau) relative to TD girls and boys (*p*=.023, *p*<.001, respectively). In addition, among boys, FC is correlated with Tau, but in opposite direction for those with ADHD (positive, *p*=.002) compared to TD (negative, *p*<.001). The last component (Tau C6) consists of subregions of the DMN (dlPFC, PCC), sensorimotor (SMA, dorsal somatomotor), and visual (lateral occipital cortex, anterior primary visual/BA17) networks with positive loadings and subregions of the cognitive control (supramarginal gyrus), DMN (superior parietal lobe), and sensorimotor (right lateralized somatomotor) networks with negative loadings, with a sex difference in TD children (*p*=.021). FC between regions in the same loading sign is significantly lower in TD boys than TD girls, while FC between regions in the opposite loading signs is higher.

### Unique components in the GNG commission error rate model (CR-C5-C7; *Figures 2h-j*)

The CAP approach identified three unique components in the GNG commission error rate model. The first component (C5) consisted of subregions of the cognitive control (IFG, dlPFC), DMN (PCC, precuneus), and sensorimotor (ACC/BA6) networks with positive loadings and subregions of the visual (secondary visual area; BA18), and dorsal attention networks with negative loadings. This component was positively associated with CR among girls with and without ADHD (*p*=.004). The second component (C6) consisted of subregions of the cognitive control (dlPFC), DMN (superior parietal lobe), and visual (posterior primary visual area; BA17) networks with positive loadings and subregions of the DMN (PCC, ACC), visual (secondary visual area; BA18), and subcortical (striatal) networks with negative loadings. This component was positively related to GAI (*p*=<.001) and negatively related to CR in girls with ADHD (*p*=.014). In addition, FC within this network was lower in boys with ADHD compared to girls with ADHD (*p*=.028) and TD boys compared to TD girls (*p*=.003). The third component (C7) consisted of subregions of the DMN (SMA, PCC) and sensorimotor (right lateralized somatomotor) networks with positive loadings and subregions of the DMN (dlPFC, ACC, superior parietal lobe), sensorimotor (left lateralized somatomotor), and auditory (opercular cortex) networks with negative loadings. This component was negatively related to GAI (*p*=.001), age (*p*=.016), and to CR in girls with ADHD (*p*=.010).

## Discussion

In this study, we present a novel statistical approach to characterize relationships between functional brain organization and cognitive function among a large cohort with ADHD and TD children. Our approach identifies higher-order networks through simultaneous modeling of multiple lower-order networks, which boosts statistical power compared to more traditional pair-wise regression approaches by substantially reducing the number of tests performed. Our results reveal complex, widespread FC patterns associated with ADHD-related sex differences involving regions of the default mode, cognitive control, somatomotor, subcortical, and visual networks. The presence and direction of these ADHD-related sex differences in FC changed when we controlled for behavioral measures of response control. These novel findings are discussed in greater detail below and placed in context of the limited existing literature examining ADHD-related sex differences.

The first network was positively related to age such that older children showed greater ITG-striatal FC and PCC-thalamic FC compared to younger children whereas PCC-striatal and ITG-thalamic FC decreased with age. To our knowledge, changes in fronto-subcortical connectivity across the 8-12 year-old age range have not been the focus of previous reports with prior studies spanning middle childhood through early adulthood (ages 8-44). Among 9 to 20 year-olds, age-related increased PCC-dorsal striatum FC and decreased FC between the ventral striatum and anterior insula and dorsal ACC have been reported.^40^ Here we did not separate dorsal or ventral striatum, suggesting an important direction for future research. Analysis of longitudinal data from 8-29 year-olds revealed age-related increases in subcortico-subcortical FC and cortico-cortical FC, whereas subcortico-cortical connections decreased with age.^41^ Our findings suggest a different pattern, with increasing subcortico-cortical FC with age, although our age-range is much smaller and different regions were examined. Finally, a cross-sectional study across 8-32 year-olds found an age-related increase in FC of the thalamus with the frontal lobe and decrease in FC of the thalamus with the temporal lobe.^42^ This is consistent with our results, providing further support for distinct age-related changes in thalamic-cortical FC for frontal and temporal regions.

The second network identified across models consists of competing contributions from FC among CC-visual-somatomotor regions with FC between CC and DMN regions and was negatively associated with GAI. Specifically, lower GAI was associated with greater FC between the (1) left central executive network, lateral occipital and ACC/BA6 regions, and (2) insula, ITG and PCC regions, and weaker FC between regions (1) and (2). Interestingly, ADHD-related sex differences were observed such that girls with ADHD displayed greater FC than did TD girls and lower FC than ADHD boys and FC within this component was positively associated with response variability only in boys with ADHD. Collectively, these findings are among the first to identify RSNs related to response variability in children with ADHD and differentially affected in girls and boys with ADHD. One previous study to our knowledge reported that atypical fronto-subcortical FC of ICA components was greatest among girls with ADHD.^18^ These findings expand upon the previous results by applying a whole-brain analytic approach to reveal more widespread alterations.

As mentioned, including response variability in the model eliminated the observed diagnostic effect in girls with ADHD. Intrasubject response variability is among the most ubiquitous findings in the ADHD neuropsychological literature,^43^ although the neural correlates of this behavioral characteristic are not well-defined. There is a lack of research examining RSNs in relation to response variability, with studies instead focusing on hemodynamic response variability during an fMRI task^44^ or associations with task-related activation.^45^ Increased response variability in ADHD is not task-specific,^46^ suggesting that identifying RSNs related to this variability may be more informative than task-specific patterns of brain activation. Our findings suggest that greater FC between executive/motor control and occipital regions and between subregions of the DMN with the insula and ITG relate to increased response variability among boys with ADHD only. Previous studies reporting that abnormal between-network connectivity is associated with ADHD symptom severity suggest that different network topology phenotypes underlie the neurobiological heterogeneity of ADHD subtypes. A similar argument could be made when examining functional network topology in relation to cognitive task performance given the established heterogeneity in cognitive deficits among individuals diagnosed with ADHD. Interestingly, FC among these regions did not differ among boys with ADHD compared to TD boys, and was not associated with response variability in TD boys, suggesting that this may be specific to heterogeneity in this cognitive process in ADHD boys rather than a more general neural correlate of ISV.

The third network we identified consists of competing contributions from FC within DMN regions with FC among DMN, CC, and visual regions and boys with ADHD showed higher FC within a DMN subregion (precuneus) with positive loadings, as well as subregions of the cognitive control (inferior frontal gyrus; IFG, insula), DMN (PCC) and visual (lateral occipital cortex) networks with negative loadings compared to TD boys. This pattern is consistent with previous findings of hyperconnectivity both within the DMN and between the DMN and other networks, particularly among boys with ADHD who are primarily, if not exclusively, examined in prior studies. Finally, this diagnostic effect in boys is not observed when controlling for GNG performance, suggesting that the FC differences observed in ADHD may contribute to response control deficits during a go/no-go task with minimal cognitive demands shown to be greatest among boys with ADHD in this age range.^25^

The fourth common network identified across models consists of competing contributions of visual-somatomotor FC and visual-DMN FC, such that boys with ADHD show hyperconnectivity between visual regions with DMN and motor networks compared to girls with ADHD. This sex difference in children with ADHD remains after accounting for response inhibition, which was greatest among boys with ADHD. However, the direction of the effect changes when response variability is accounted for in the Tau model such that girls with ADHD show increased FC in this network compared to TD boys demonstrating similar response variability. Finally, FC within this network is negatively associated with both tau and commission error rate in boys with ADHD, suggesting a possible sex-specific neural correlate, as discussed above. This pattern of findings suggests that visual-somatomotor and visual-DMN FC is differentially affected in girls and boys with ADHD, with boys generally showing greater anomalies and associations with response control.

The unique components of interest from the Tau model (C5) showed a diagnostic difference in girls and boys with ADHD and associations with tau in boys only, while the unique component of interest from the CR model (C5-C7) showed a sex difference in children with ADHD (C6) and associations with CR in girls with ADHD (C5-C7). Our behavioral findings are consistent with prior reports of ADHD-related sex differences in response inhibition, such that girls with ADHD show intact response inhibition compared to TD girls when cognitive demands are minimal, whereas boys with ADHD make significantly more response inhibition errors.^25^ However, within the group of girls with ADHD, individual differences in response inhibition are important to consider, with some girls with ADHD showing impairments in this cognitive process. The results from the three unique components of the CR model suggest that individual variability among subregions of the cognitive control, DMN, sensorimotor, visual, and dorsal attention networks relates to response inhibition among girls with ADHD.

In conclusion, using a novel statistical approach to identify a common linear projection of multiple time courses across subjects such that variations in functional connectivity defined by the projection can be explained by the covariates of interest, we found a complex pattern of ADHD-related sex differences that would be difficult to capture using traditional pairwise regression approaches. These findings add to the growing literature suggesting that the neurobiological basis for ADHD may differ among girls and boys.^18,28,47,48^ This is also important in advancing our understanding of ADHD-related sex differences in cognitive functions, which may suggest different etiological pathways for girls and boys with ADHD that may ultimately inform prevention and intervention approaches. Future studies oversampling girls with ADHD, as we have done, will be important for replicating these findings and establishing the utility of this novel approach to analysis of intrinsic functional network topology in relation to ADHD and associated cognitive functions.

## Materials and Methods

### Participants

Participants include 217 8-12 year-old children with either a diagnosis of ADHD (n=115, 29 girls) or TD controls (n= 102, 35 girls). Participants were recruited through local schools, community-wide advertisement, volunteer organizations, medical institutions, and word of mouth. This study was approved by the local Institutional Review Board. After providing a complete study description to the participants, oral informed consent was obtained from a parent/guardian followed by an initial phone screening. Children with a history of intellectual disability, learning disability, seizures, traumatic brain injury, or other neurological illnesses were excluded. Participants were determined to be eligible for inclusion in either the ADHD group or the TD group based upon review of standardized rating scales and diagnostic interview (see **Supplementary Materials** for details of the diagnostic procedure). Eligible participants and their parents attended two laboratory sessions. At the initial visit, written informed consent and assent were obtained from the parent/guardian and the child and intellectual ability was assessed. Children taking psychotropic medications other than stimulants were excluded from participation and children taking stimulants were asked to withhold medication the day prior to and day of testing. Basic demographic information is provided in **Table 2**.

**Table 2.**
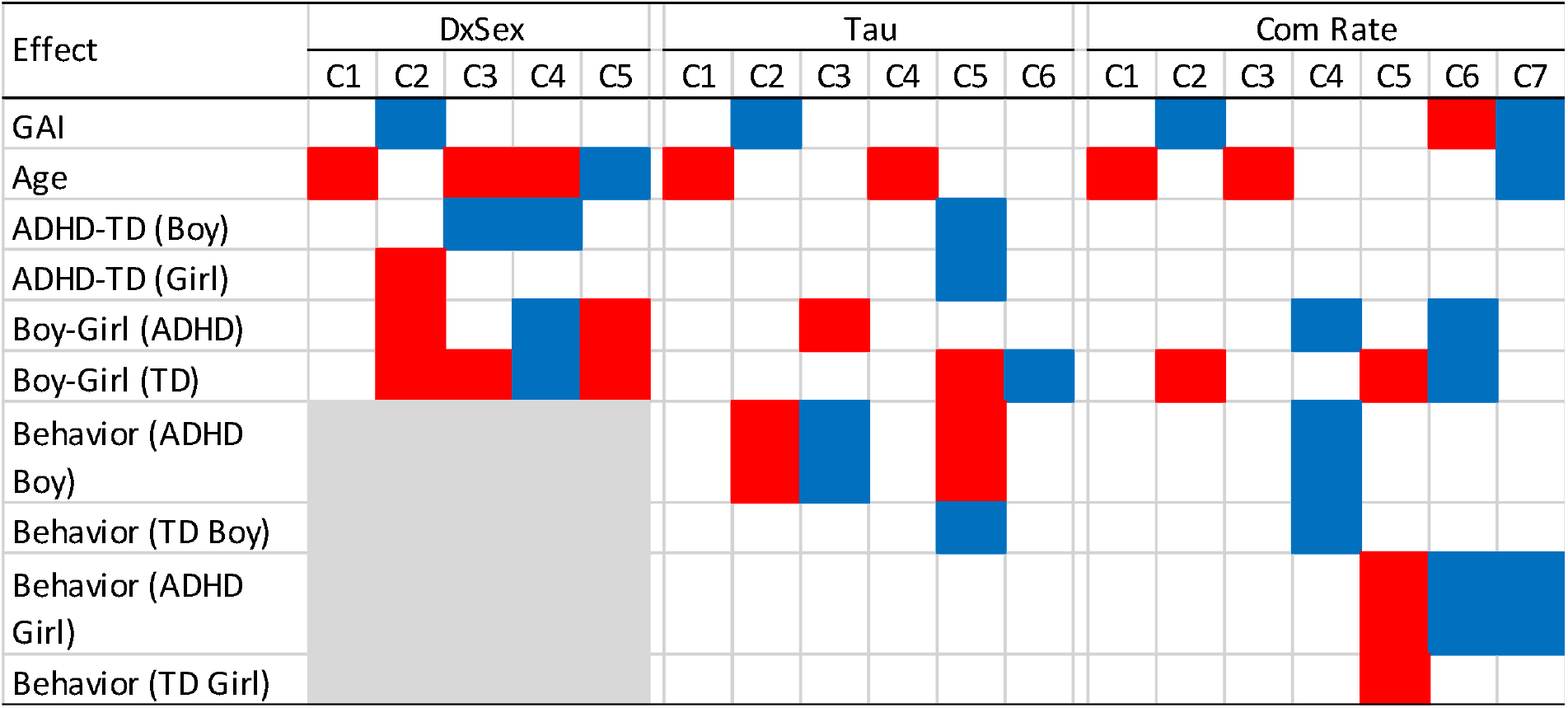
Significant effect of the components estimated from the CAP model. The red color denotes a positive effect and blue for negative. DxSex: the behavior free model; Tau: model including Go/No-Go Tau as one of the predictors; Com Rate: model including Go/No-Go commission error rate as one of the predictors.

### Behavioral Go-No/Go (GNG) data

Participants completed a GNG task, from which we have previously published findings from a subset of this sample examining behavioral performance and relations with anatomical imaging data.^18,24–31^ The GNG task was programmed in Presentation (Neurobehavioral Systems, Albany, CA, USA). Participants were seated in front of a computer monitor with a keyboard. The task stimuli consisted of presentation of a single green spaceship for “Go” trials (80%) and a red spaceship for “No-Go” trials (20%) presented for 300 ms with an interstimulus interval of 2000 ms during which a fixation cross appeared across 240 trials. Participants were instructed to push the spacebar with their index finger as quickly as possible in response to green spaceships and withhold responding to red spaceships. Response inhibition was quantified as commission error rate (CR), calculated as the proportion of no-go trials on which participants incorrectly responded. Response variability was quantified using the ex-Gaussian indicator, tau, representing infrequent, slow responses contributing to the exponential component of the reaction time (RT) distribution.^32^

### Resting-state fMRI data

All children completed a mock scan to acclimate to the scanning environment and then rs-fMRI was acquired on a 3.0 T Philips scanner (see **Supplementary Materials** for acquisition and preprocessing details and prior publications from a subset of this sample examining functional connectivity without relating this to GNG behavioral performance).^18,33,34^ We decomposed the rs-fMRI data into temporally coherent networks using the Group ICA of fMRI Toolbox (GIFT: http://icatb.sourceforget.net; Medical Image Analysis Lab, Albuquerque, New Mexico).^23,35^ We chose ICA^36^ rather than seed-based approaches because of its effectiveness at separating signal from noise,^37^ its increased sensitivity to detecting individual differences,^38^ and its ability to identify resting state networks (RSNs) without defining a seed region by effectively clustering voxels with similar timecourses. An additional benefit of this data-driven grouping of functionally related voxels is that it reduces the size of the covariance matrix used in the CAP model and thereby allows for whole-brain analysis. Following group ICA with backward reconstruction (see **Supplementary Materials**), we identified relevant RSNs by comparing the spatial distribution of each of the group-level, aggregate ICs to a publicly available set of unthresholded IC t-maps that have been classified as RSNs by a group of experts and organized into seven large functional groups: visual (Vis), auditory (Aud), somatomotor (SM), default mode (DMN), cognitive control (CC), sub-cortical (SC) and cerebellar (Cb) networks.^39^ As an added precaution against the contamination of brain networks from persistent motion-related variance, we regressed motion covariates from IC timecourses associated with components identified as representing signal before estimating between-component functional connectivity (FC).^39^

### Covariate Assisted Principal (CAP) Regression on RSN Components

Details of the CAP regression on the signal components identified by group ICA described above is provided in the **Supplementary Materials**.^21^ The CAP method identifies a linear projection of the covariance matrices such that between-subject variability in FC is most strongly associated with the covariates of interest. Assuming the IC time courses are standardized to have identical variance, the CAP regression models the association between FC and the covariates. This association depends on both the sign of the model coefficient and the sign of the loading products. For a positive coefficient estimate, FC between two ICs with the same loading sign (then the product is positive) is positively associated with the corresponding covariate; while FC for two ICs with the opposite signs (then the product is negative) is negatively associated with the covariate.

### Analysis

We modeled three sets of covariates. First, we tested for effects of an ADHD diagnosis, sex and their interaction, as well as age and intellectual reasoning ability (general ability index; GAI) in the CAP regression model. In subsequent models, we examined associations with behavioral measures of response control, GNG Tau (log-transformed) and CR. These models included ADHD diagnosis, sex, the behavioral measure of interest and all possible interactions among these three variables.

## Acknowledgements

This research was supported by the National Institutes of Health awarded to Dr. Stewart Mostofsky (R01MH078160 and R01MH085328), Dr. Keri Rosch (K23MH101322), Dr. Mary Beth Nebel (K01MH109766), and Dr. Karen Seymour (K23MH107734) and by the Intellectual and Developmental Disabilities Research Center at Kennedy Krieger Institute and The Johns Hopkins University under award number NIH P50HD103538.

