## Supplemental File for "Beyond massive univariate tests: Covariance regression reveals complex patterns of functional connectivity related to attention-deficit/hyperactivity disorder, age, sex, and response control"

**Supplementary Materials**

***Diagnostic procedures.*** Intellectual ability was assessed using the Wechsler Intelligence Scale for Children, Fourth Edition (WISC-IV)^1^ or Fifth Edition (WISC-V).^2^ Participants with general ability index (GAI) scores, a measure of intellectual reasoning ability that does not factor in working memory and processing speed scores, below 80 were excluded. To screen for reading disorders, children were administered the Word Reading subtest from the Wechsler Individual Achievement Test, Second Edition (WIAT-II)^3^ or Third Edition (WIAT-III)^4^ and were excluded for standard scores below 85. Diagnostic status was established through administration of either the Diagnostic Interview for Children and Adolescents, Fourth Edition (DICA-IV)^5^ or the Kiddie Schedule for Affective Disorders and Schizophrenia for School Aged Children Present Lifetime version (KSADS-PL).^6^ Children meeting criteria for diagnosis of conduct, mood, generalized anxiety, separation anxiety or obsessive–compulsive disorders on either interview were excluded whereas a comorbid diagnosis of oppositional defiant disorder (ODD) was permitted. Parents and teachers (when available) also completed the Conners Parent and Teacher Rating Scales-Revised Long Version or the Conners-3 (CPRS and CTRS)^7,8^ and the ADHD Rating Scale-IV (ADHD-RS), home and school versions.^9^ A diagnosis of ADHD was confirmed by a child neurologist or psychologist based on the diagnostic interview, which considered information provided by the parent about functioning at school, in addition to onset, course, duration, and frequency of symptoms, and parent/teacher rating scales (i.e., T-scores ≥ 65 or ≥ 6 symptoms endorsed on at least one rating scale). Inclusion in the TD group required scores below clinical cutoffs (i.e., T-scores ≤ 60 and ≤ 4 symptoms endorsed on all parent/teacher rating scales.

***Resting-state fMRI data acquisition****.* Resting state fMRI (rs-fMRI) was acquired on a 3.0 T Philips scanner using a single-shot, partially parallel, gradient-recalled echo planar sequence with sensitivity encoding and an ascending slice order (repetition time [TR]/echo time [TE] = 2500/30ms, flip angle = 75^o^, sensitivity encoding acceleration factor of 2, 47 3-mm axial slices with no slice gap, in-plane resolution of 3.05×3.15 mm [84×81 voxels], duration = 5 min 20 sec – 6 min 30 s). Participants were instructed to relax, fixate on a cross-hair, and remain as still as possible.

***Preprocessing of fMRI data****.* Functional data were preprocessed using SPM12 (Wellcome Trust Centre for Neuroimaging, London, United Kingdom) and custom MATLAB (The Mathworks, Inc., Natick, Massachusetts) code. rs-fMRI scans were slice-time adjusted using the slice acquired in the middle of the TR as a reference, and rigid body realignment parameters were estimated to adjust for motion. The volume collected in the middle of the scan was spatially normalized using the Montreal Neurological Institute (MNI) EPI template.^12^ The estimated rigid body and nonlinear spatial transformations were applied to the functional data together, producing 2-mm isotropic voxels in MNI space. Linear trends were removed, and the data were spatially smoothed using a Gaussian filter (6-mm full width at half maximum kernel). Participants were excluded for between-volume translational movements >3 mm or rotational movements >3 degrees. Mean framewise displacement (FD) was calculated using the realignment estimates.^10^ A significant challenge in the ADHD neuroimaging literature is accounting for artifacts introduced by motion during the scan, which have been shown to impact FC metrics and usually differ between groups with and without psychopathology. Various approaches to minimizing the effects of motion on FC have been proposed, including matching clinical and control groups on framewise displacement (FD, a measure of head motion between consecutive fMRI volumes),^11^ covarying for FD, and removing volumes contaminated by motion within an fMRI scan. One disadvantage of these approaches is that variance associated with ADHD symptomatology and cognitive deficits may not be independent from variance associated with motion; excluding children with ADHD who move more during the scan may bias the sample towards less severely affected children. We chose to not match the groups on mean FD, given evidence that head motion is correlated with ADHD symptomatology and that head motion and ADHD may have similar genetic loadings.^12^ A growing body of literature has demonstrated the effectiveness of ICA-based strategies for identifying and removing motion-related variation in fMRI data.^13,14^ Here, we utilized spatial ICA to isolate motion-related sources from functional network sources. As an added precaution against the contamination of brain networks from persistent motion-related variance, we regressed motion covariates from ICA-based brain network timecourses before estimating between network functional connectivity.^15^

***ICA with Backward Reconstruction****.* We used an information theoretic approach to dimension estimation^16^ and chose the number of independent components (ICs) for the group to be the maximum dimension estimate across participants, 65. Prior to ICA, each participant’s preprocessed data were variance normalized on a voxelwise basis and reduced to 100 principle components (PCs) using principal component analysis (PCA). Participant-specific PCs were temporally concatenated and a second PCA was used to reduce the aggregate data set to the maximum dimension estimated, 65 (defined above) using multi-power iteration.^17^ ICA was repeated on the group-level PCs 100 times using the Infomax algorithm^18^ and the ICASSO toolbox^19^ with randomized initial conditions in GIFT to ensure stable ICs. Participant-specific spatial maps (SMs) and timecourses (TCs) were generated from the aggregate IC decomposition using a method based on PCA compression and projection.^20^ The SMs represent the spatial topography of each component within the brain while the TCs represent the intrinsic level of engagement of each component over time.

***Covariate Assisted Principal Regression on RSN Components***

We implemented covariate assisted principal (CAP) regression on the signal components identified by group ICA described above.^21^ For subject
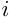
, let
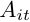
 denote the
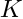
 IC time courses at time
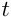
 for
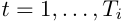
. The CAP approach assumes that there exist orthogonal linear projections
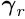
 for
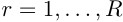
 (
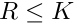
), such that in the projection space, the data variation satisfies the following log-linear model:

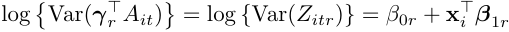
,

where
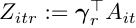
;
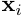
 is the vector of covariates of interest;
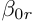
 and
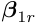
 are model coefficients;
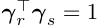
 if
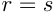
 and 0 otherwise. The projections and model coefficients can be estimated by maximizing the likelihood function assuming
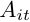
 is normally distributed with mean zero and covariance matrix
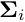
 (for
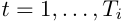
 and
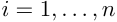
). The number of projections,
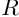
, is determined based on the level of deviation from diagonality.^22^ To draw inference about the model coefficients, 95% confidence intervals were acquired from 500 bootstrap samples fixing the estimated linear projection. The method was implemented using the R package cap available on CRAN.

CAP components were reconstructed in voxel space to form brain maps representing orthogonal groups of signal ICs associated with the variables of interest. Let
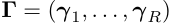
 denote the orthonormal projection matrix estimated by CAP regression. Each row of
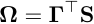
 then represents the newly constructed CAP brain map, where
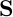
 is the spatial maps obtained from the group ICA.

The CAP method identifies a linear projection of the covariance matrices such that between-subject variability in FC is most strongly associated with the covariates of interest. Let
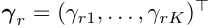
 and
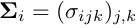
 for
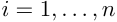
, then

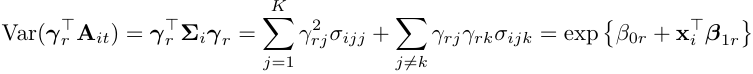
.

Assuming the IC time courses are standardized to have identical variance (for example,
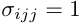
 for any
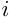
 and
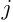
), the CAP regression models the association between FC and the covariates. This association depends on both the sign of the
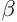
 coefficient and the sign of the loading products. For a positive
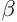
 estimate, FC between two ICs with the same loading sign (then the product is positive) is positively associated with the corresponding covariate; while FC for two ICs with the opposite signs (then the product is negative) is negatively associated with the covariate.

**Supplementary Figure S1.** River plot of IC loadings of the components identified by the CAP method for the (a) DxSex (b) Tau and (c) ComRate models. The IC indexes are color coded by the functional module.

(a)

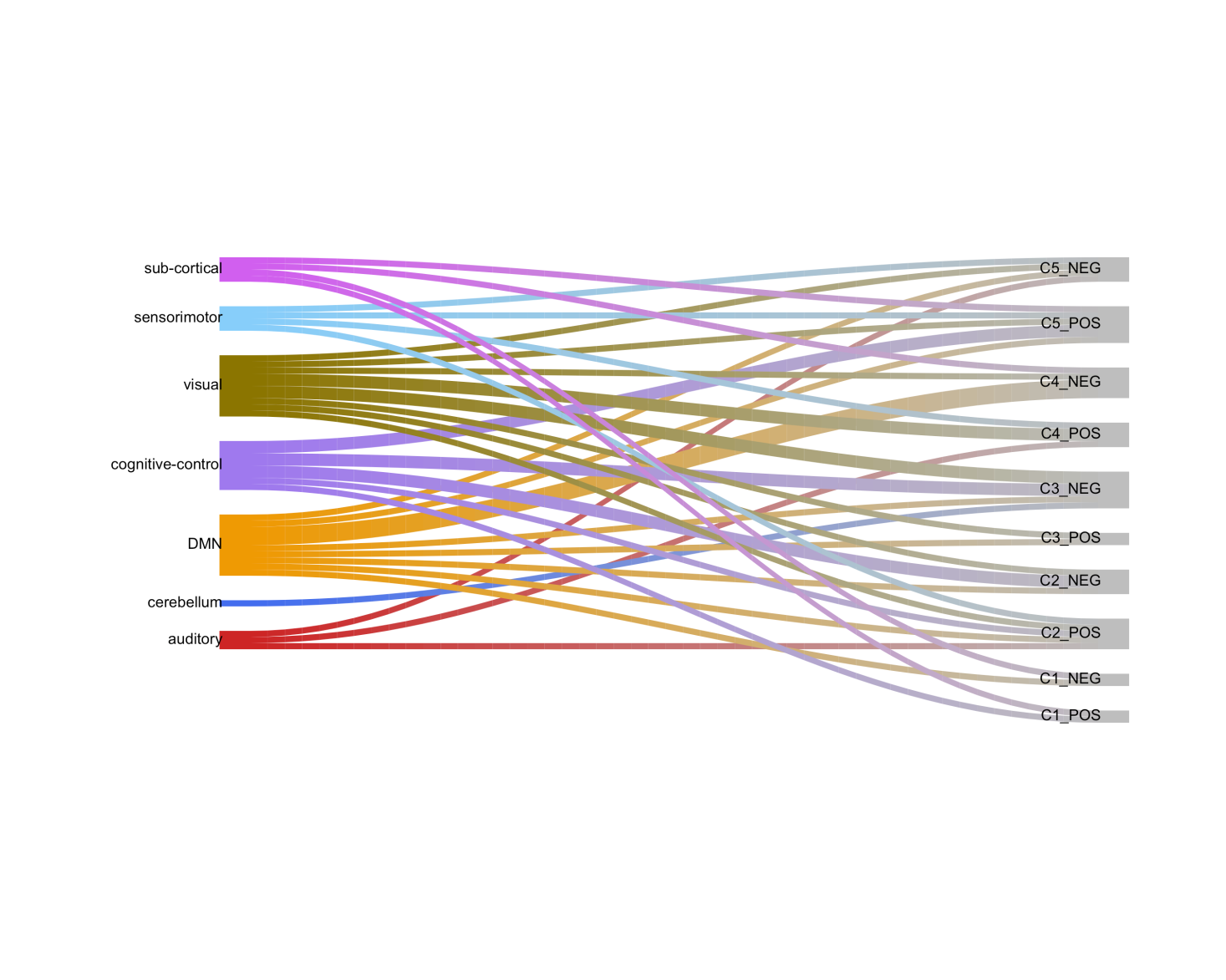

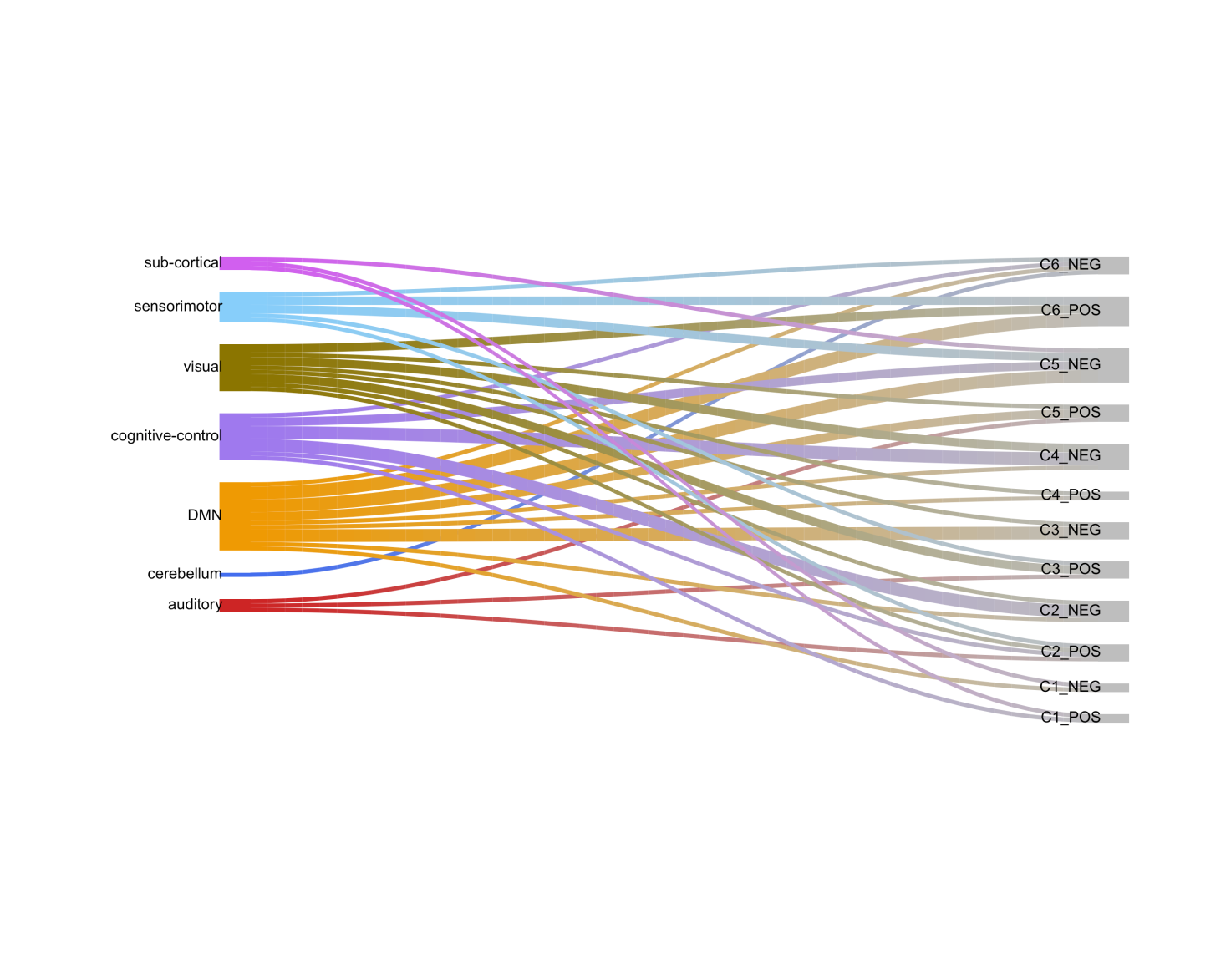

(c)

(b)

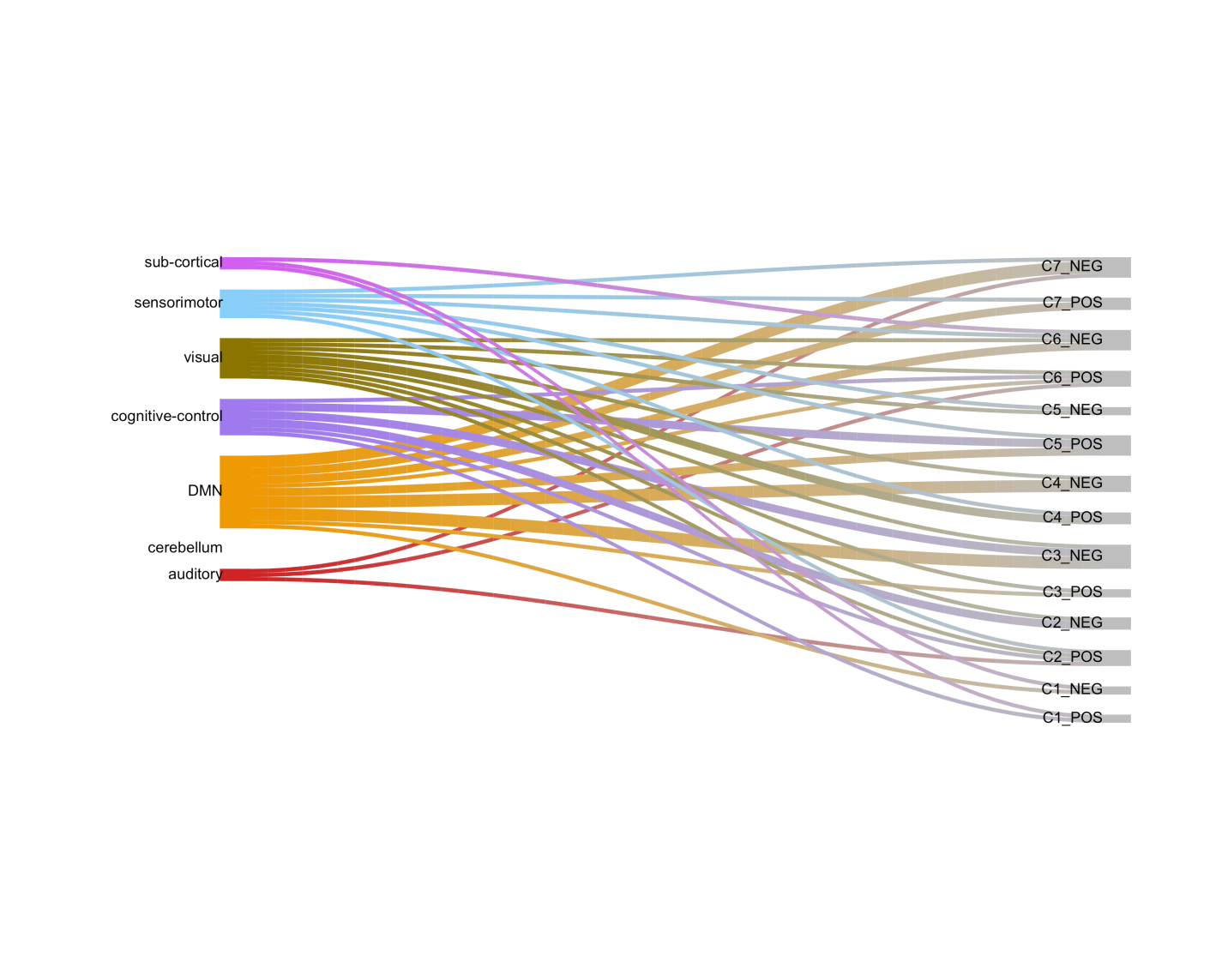

**Supplementary Table 1.** Coefficients and 95% confidence intervals obtained from 500 bootstrap samples in the three models.

(a) The Dx-Sex model

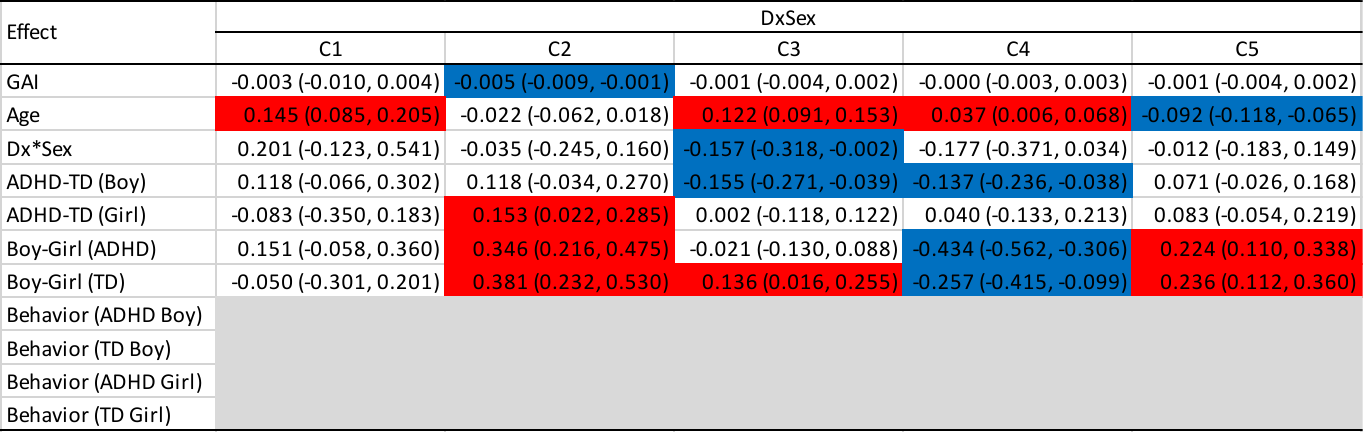

(b) The GNG Tau model

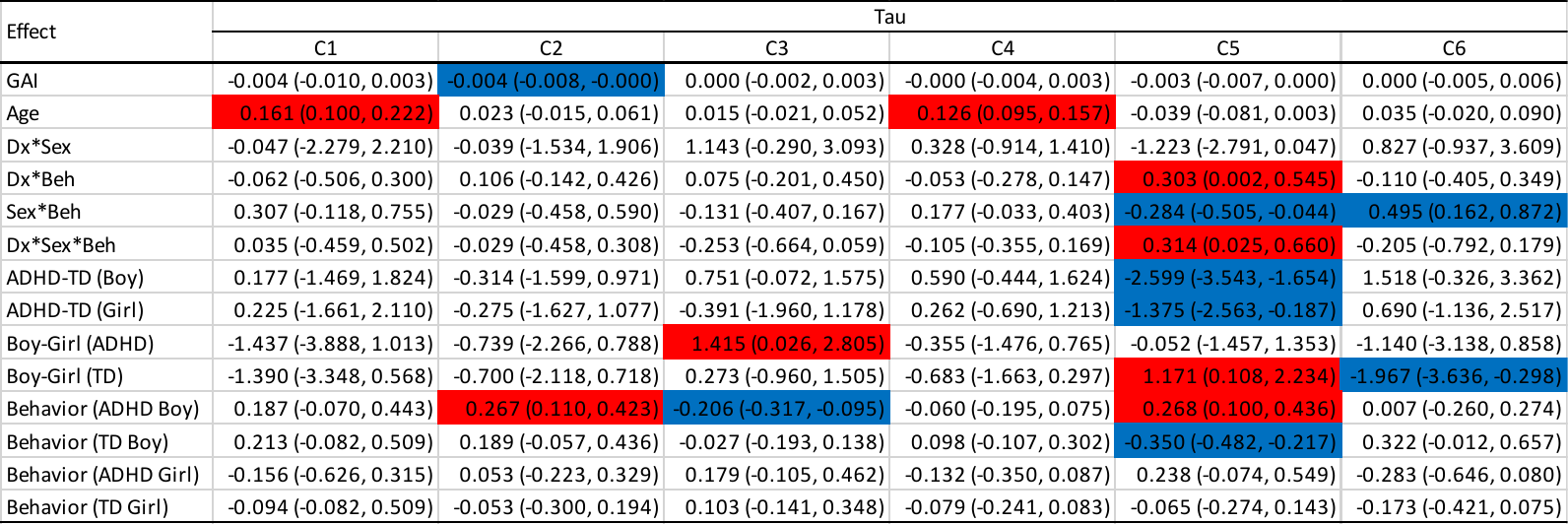

(c) The GNG Commission Error Rate (CR) model

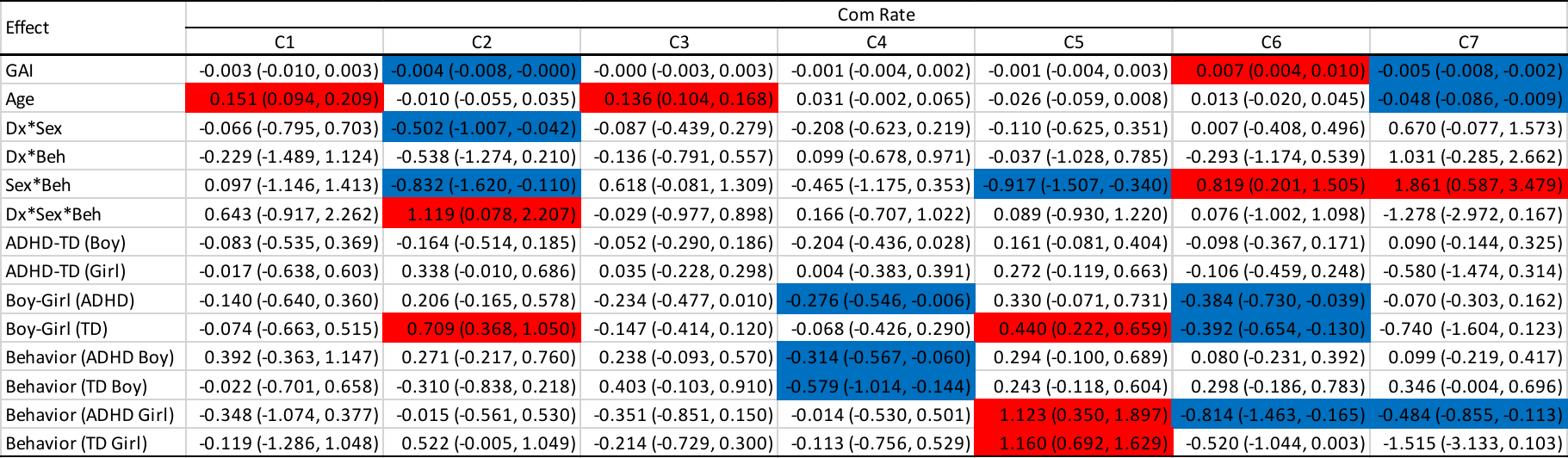
